# Comparing Models of Information Transfer in the Structural Brain Network and Their Relationship to Functional Connectivity: Diffusion Versus Shortest Path Routing

**DOI:** 10.1101/2022.11.11.515707

**Authors:** Josh Neudorf, Shaylyn Kress, Ron Borowsky

## Abstract

The relationship between structural and functional connectivity in the human brain is a core question in network neuroscience, and a topic of paramount importance to our ability to meaningfully describe and predict functional outcomes. Graph theory has been used to produce measures based on the structural connectivity network that are related to functional connectivity. These measures are commonly based on either the shortest path routing model or the diffusion model, which carry distinct assumptions about how information is transferred through the network. Unlike shortest path routing, which assumes the most efficient path is always known, the diffusion model makes no such assumption, and lets information diffuse in parallel based on the number of connections to other regions. Past research has also developed hybrid measures that use concepts from both models, which have better predicted the functional connectivity from structural connectivity than shortest path length alone. We examined the extent to which each of these models can account for the structure-function relationship of interest using graph theory measures that are exclusively based on each model. This analysis was performed on multiple parcellations of the Human Connectome Project using multiple approaches, which all converged on the same finding. We found that the diffusion model accounts for much more variance in functional connectivity than the shortest path routing model, suggesting that the diffusion model is better suited to describing the structure-function relationship in the human brain at the macroscale.

---

Graph theory analyses of structural brain connectivity have been vital to providing breakthroughs in our understanding of how the underlying structure of the brain can influence the patterns of coordinated functional activity (see Avena-Koenigsberger et al., 2018 for a review; see also Goñi et al., 2014; Neudorf, Ekstrand, et al., 2020; Neudorf et al., 2022; Neudorf, Kress, et al., 2020). Defining this relationship between structural and functional connectivity using advanced techniques including graph theory has recently been highlighted as an important frontier in neuroscience (Suárez et al., 2020). When it comes to choosing graph theory measures of connectivity, important assumptions must be made about how information is transferred through the structural network, and the effectiveness of these measures for predicting functional connectivity is dependent on the accuracy of these assumptions about the human brain. Two primary graph theory models of information transfer in the brain include shortest path routing and diffusion. The shortest path routing model relies on the calculation of the shortest path to the destination region. This model is straightforward to calculate and underlies many useful graph theory measures that have been helpful in describing brain networks and networks in general (e.g., characteristic path length, Watts & Strogatz, 1998; global efficiency as an indicator of small-worldness, Latora & Marchiori, 2001; nodal and local efficiency, Latora & Marchiori, 2001; van den Heuvel & Sporns, 2013; etc.). One problem with the shortest path routing model when it comes to brain networks is that it assumes each region has whole-brain level knowledge about the most efficient path to use (Avena-Koenigsberger et al., 2019; Seguin et al., 2018, 2022; Zamani Esfahlani et al., 2022).

An alternative graph theory model has been proposed that does not assume whole-brain knowledge about the shortest path, but instead assumes that information diffuses along random paths in the network influenced by the relative weighting of each path. Under these model assumptions, information propagates through the network as a “random walker” that is constrained by the structural architecture. Furthermore, information can be transferred in parallel, whereas shortest path routing describes information traveling along a single path to the destination (Fornito et al., 2016a).

Novel graph theory metrics combining both diffusion and shortest path routing models have been developed for use in brain research and applied to the task of predicting functional connectivity from the underlying structural connectivity (Goñi et al., 2014). *Search information* was developed as a measure of how many distractor paths may lead a random walker away from the shortest path, while *path transitivity* measures how likely a random walker on a detour will end up back on the shortest path. While these measures were more successful than shortest path length alone at predicting functional connectivity from structural connectivity, there are other graph theory measures of connectivity that consider the full range of possible paths based on a diffusion model, rather than hinging on what happens around the shortest path during information transfer. One such measure of diffusion efficiency is the *mean first passage time* (Wang & Pei, 2008), which calculates the number of steps it takes a random walker on average to travel from region A to region B. This measure has been used to show that biological brain networks typically display a balance between diffusion efficiency and global efficiency (sensitive to shortest path length; Goñi et al., 2013). Another measure relying on the diffusion model of information transfer is *communicability* (Estrada & Hatano, 2008), which takes into consideration all possible walks from region A to region B. Walks with less edges, *n*, are weighted much higher than those with more, with walks weighted by the factor 1/*n*!. *Communicability* is described as reflecting the capacity for a network to transfer information in parallel assuming a diffusion model of information transfer (Fornito et al., 2016). This measure has been useful in distinguishing patients from controls, including stroke (Crofts et al., 2011) and multiple sclerosis (Li et al., 2013).

Considering past success with the hybrid measures combining the diffusion and shortest path routing models of information transfer (Goñi et al., 2014), this research will apply the exclusively diffusion-based measures of mean first passage time and communicability as well as the shortest path routing measure of shortest path length to structural connectivity, and the results will be used to predict functional connectivity to determine to what extent these measures are able to account for variance in functional connectivity. Crucially, this research will extend past research that has examined the ability of multiple graph theory communication measures to predict functional connectivity from structural connectivity (Betzel et al., 2022; Vázquez-Rodríguez et al., 2019; Zamani Esfahlani et al., 2022) and benchmarking the ability for different communication measures to predict functional connectivity (Seguin et al., 2018, 2020, 2022), by directly comparing two commonly used models (diffusion and shortest path routing) using multiple linear regression analyses, partial least squares regression, and principal components analysis to determine which graph theory model is most important in this relationship. Research suggests that brain networks (at both the macroscale and microscale) typically demonstrate a balance of diffusion efficiency and global efficiency (Goñi et al., 2013), while also suggesting that this balance may lean more towards dominance of diffusion efficiency in human brains, in which case we expect that the diffusion measures examined here will be more relevant than shortest path length to the structure-function relationship in the brain.

## Methods

### Dataset

MRI data for 998 subjects from the Human Connectome Project (HCP) were used including DTI and rsfMRI (Van Essen et al., 2013). We used the preprocessed version of the rsfMRI data. This data has been preprocessed using FSL FIX (Salimi-Khorshidi et al., 2014). The DTI data used was also preprocessed. The HCP pipelines for preprocessing are described by Glasser et al. (2013). The Automated Anatomical Labelling 90 region atlas (AAL; Tzourio-Mazoyer et al., 2002) was used as well as the Brainnetome 246 region atlas (Fan et al., 2016). Activation at each rsfMRI acquisition was used to calculate the mean activation for the atlas regions. The rsfMRI sessions were standardized using a z-score for the regions for each session separately. The activation in these regions was then submitted to bandpass filtering (separately for each session) allowing only frequencies within 0.01 Hz and 0.1 Hz (see Hallquist et al., 2013).

### Connectivity measures

To calculate the functional connectivity measures for each combination of regions, we calculated the Pearson correlation coefficient using all of the 4800 acquisitions. To calculate the structural connectivity measures, DSI Studio (http://dsi-studio.labsolver.org) was used with quantitative anisotropy (Yeh et al., 2013) as the termination index to calculate the streamline count. Generalized Q-sampling (Yeh et al., 2010) was used, and tracking used 1 million fibers, 75 degrees maximum angular deviation, and a 20 mm minimum and 500mm maximum fiber length. To calculate the structural connectivity matrix containing the number of streamlines for each cell, a whole brain seed was used. The connectivity values for structural and functional connectivity were averaged using the mean for all subjects. The weighted structural connectivity density (the sum of connection weights divided by the total possible connection weights, where each weight has a maximum of 1.0) was .016 for the AAL atlas and .004 for the Brainnetome atlas.

Graph theory structural connectivity measures of mean first passage time (Wang & Pei, 2008) and communicability (Estrada & Hatano, 2008) were calculated as diffusion model measures (also discussed in Fornito et al., 2016). *Mean first passage time* was calculated as

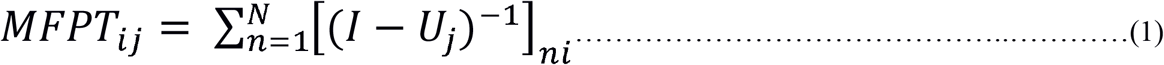

where *I* is the identity matrix, *i* is the starting node, *j* is the destination node, and *N* is the number of regions in the network, and

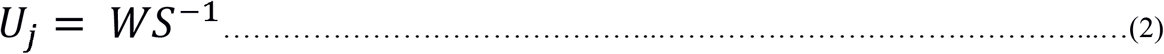

but with the *j*th row set to zero so that a random walker is unable to enter *j. W* is the weighted structural connectivity adjacency matrix, and

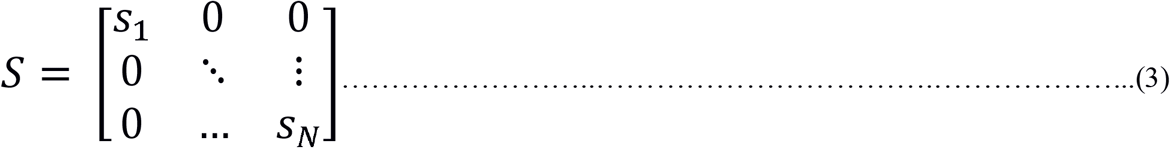

with *S*_*n*_ representing the strength (weighted number of connections) of region *n. Communicability* was calculated as

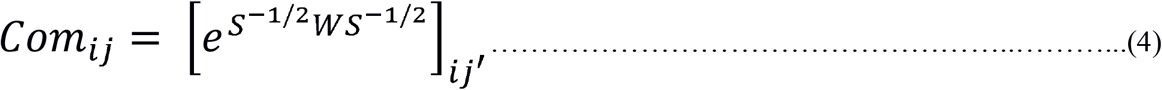

where 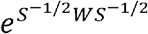 is the matrix exponential of 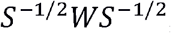, the reduced structural connectivity adjacency matrix (see Crofts et al., 2011). Long walks are weighted more weakly (by a factor of *n!* where *n* is the number of steps) in this formula as the series expansion equates to

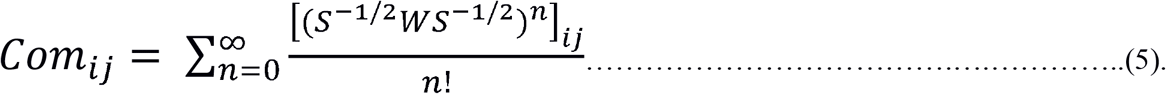

*Shortest path length* was calculated as the shortest path routing model measure using the NetworkX python library (Hagberg et al., 2008; function *shortest_path_length*, using the Dijkstra algorithm described by Dijkstra, 1959, and given the inverse value of structural connectivity edges so that the edges represent resistance in the network).

Permutation testing was performed using 500 null models created from the structural connectivity following the generalized Maslov-Sneppen (Maslov & Sneppen, 2002) rewiring algorithm developed by Rubinov & Sporns (2011) for use with weighted networks to control for node strength and degree while randomizing the connection weights. Permutation *p*-values were calculated as the number of null models resulting in the same or better variance accounted for in the models as a proportion of the total number of null models.

## Results

### AAL

#### Linear regression models

Linear regression models were computed for each log transformed independent variable (mean first passage time, communicability, and shortest path length) with functional connectivity as the dependent variable using the *lm* function from the *lme4* library (Bates et al., 2015) in *R* (R Core Team, 2018). Mean first passage time demonstrated an inverse relationship with functional connectivity, whereby a high mean first passage time was associated with poorer functional connectivity, as expected, *R*(4003) = -.376, *p* < .001 (null model permutation *p* = .044; see Figure 1A). Communicability demonstrated a positive relationship with functional connectivity, whereby high communicability was associated with better functional connectivity, as expected, *R*(4003) = .316, *p* < .001 (null model permutation *p* = .014; see Figure 1B). Shortest path length demonstrated an inverse relationship with functional connectivity, whereby a high shortest path length was associated with poorer functional connectivity, as expected, *R*(4003) = -.379, *p* < .001 (null model permutation *p* < .002; see Figure 1C). The magnitude of the structure-function relationship for each of these measures was relatively comparable, so additional multiple linear regression approaches were also taken to determine which measures are primarily driving the relationship between structural and functional connectivity.

**Figure 1.**
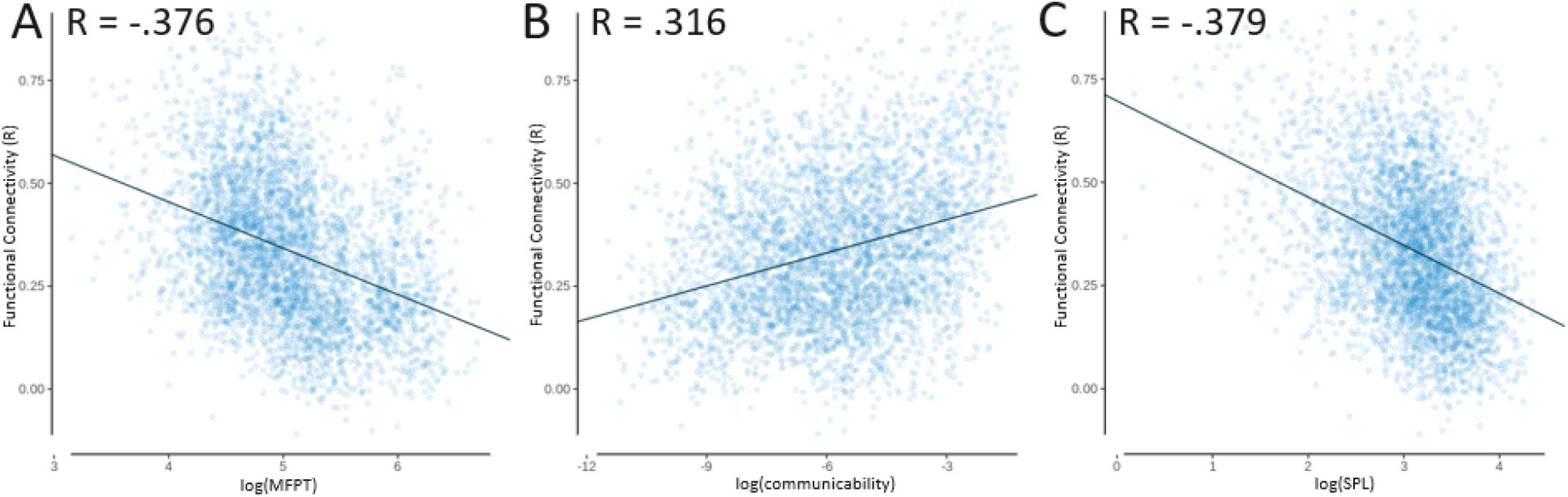
AAL linear regression models with functional connectivity as the independent variable and dependent variables of: **(A)** mean first passage time (MFPT; log transformed), *R*_*adj*_ = -.376; **(B)** communicability (COM; log transformed), *R*_*adj*_ = .316; **(C)** and shortest path length (SPL; log transformed), *R*_*adj*_ = -.379.

#### Multiple Linear Regression Models

Multiple linear regression models were then investigated starting with mean first passage time and shortest path length included in the model as independent variables, with functional connectivity as the dependent variable. These models were again calculated in *R* using the *lm* function from the *lme4* library, as well as *spcor* from the *ppcor* library to calculate the semi-partial correlation (Kim, 2015) and *vif* from the *car* library to calculate the variance inflation factor (Fox & Weisberg, 2019). Prior to this, the correlation matrix of these measures was examined, which indicated that there were no extreme correlations (e.g., greater than .9) between the independent variables, with the highest value being *R* = .841 between mean first passage time and shortest path length (see Table 1). This correlation is theoretically interesting though, as it indicates there is a high level of redundancy between mean first passage time and shortest path length, suggesting that information in the diffusion model naturally follows paths that are similarly efficient when compared to the shortest path. This potential for decentralized information transfer strategies to take advantage of shortest paths in the network has been noted in past research (Avena-Koenigsberger et al., 2017; Goñi et al., 2014; Seguin et al., 2018; Vézquez-Rodríguez et al., 2020). As seen in Table 2, mean first passage time and shortest path length both produced significant effects, with shortest path length having a slightly larger semi-partial correlation (see Figure 2A for predicted vs. empirical functional connectivity). However, when adding communicability to the model as seen in Table 3, the overall variance accounted for increased, and the semi-partial correlation of shortest path length was greatly reduced (though still significant), while the diffusion-based measures of mean first passage time and communicability had a much larger combined magnitude of semi-partial correlation. This model accounted for more variance than any of the measures independently (*R*^2^ = .165; see Figure 2B for predicted vs. empirical functional connectivity). It should be noted that the variance inflation factor (VIF) for shortest path length in model 2 was greater than 5 (*VIF* = 5.567), indicating that multicollinearity between the independent variables may have affected the variance of the shortest path length coefficient. In order to address this, we also examined these variables using partial least squares regression, which is robust against multicollinearity.

**Table 1.**
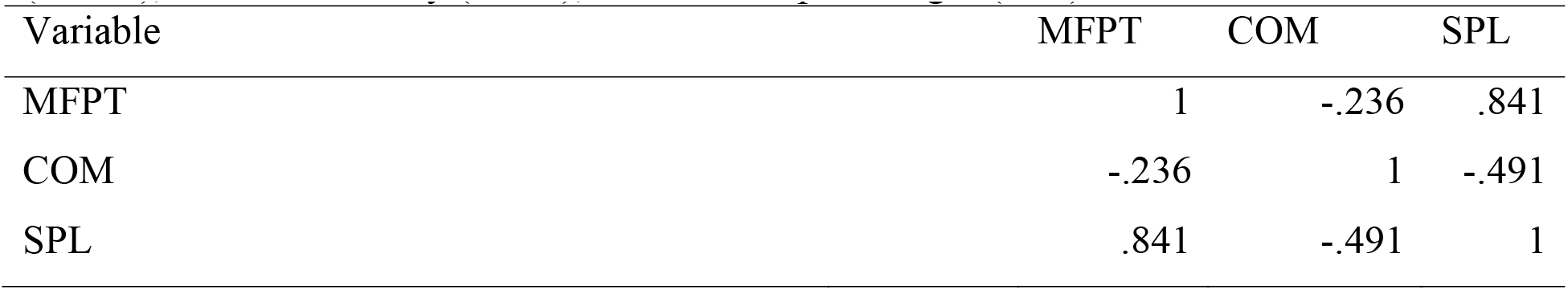
AAL independent variable correlation matrix. Variables include mean first passage time (MFPT), communicability (COM), and shortest path length (SPL).

**Table 2.** AAL multiple linear model 1, with dependent variable functional connectivity. *R*^*2*^ = .157, *R*_*adj*_^2^ = .157 (null model permutation *p* = .012). Abbreviations include semi-partial correlation (SPC) and variance inflation factor (VIF).

| Effects | SPC | Estimate | Std. Error | $t$-value | $p$-value | VIF |
| --- | --- | --- | --- | --- | --- | --- |
| Intercept |  | .845 | .023 | 36.577 | <.001 |  |
| log(MFPT) | -.117 | -.060 | .007 | -8.051 | <.001 | 2.927 |
| log(SPL) | -.127 | -.067 | .008 | -8.745 | <.001 | 2.927 |

**Table 3.** AAL multiple linear model 2, with dependent variable functional connectivity. *R*^*2*^ = .165, *R*_*adj*_^2^ = .165 (null model permutation *p* = .016). Abbreviations include semi-partial correlation (SPC) and variance inflation factor (VIF).

| Effects | SPC | Estimate | Std. Error | $t$-value | $p$-value | VIF |
| --- | --- | --- | --- | --- | --- | --- |
| Intercept |  | .851 | .023 | 36.960 | <.001 |  |
| log(MFPT) | -.138 | -.075 | .008 | -9.592 | <.001 | 3.229 |
| log(COM) | .089 | .012 | .002 | 6.184 | <.001 | 2.493 |
| log(SPL) | -.031 | -.022 | .010 | -2.112 | .035 | 5.567 |

#### Partial least squares regression

A partial least squares regression analysis was conducted with a dependent variable of functional connectivity and independent variables of mean first passage time, communicability, and shortest path length, using the *plsr* function from the *pls* library in *R* (Mevik & Wehrens, 2007). The independent variables were log transformed and standardized to have a mean of 0 and standard deviation of 1. To validate the model and check for overfitting a k-fold cross-validation scheme was used with 10 folds. The number of components to include was decided when additional components no longer substantially decreased the root mean squared error of prediction. With 2 components included, the root mean squared error of prediction reached its minimum of .161, so 2 components were used. Cross-validation determined that the model was able to account for 16.3% (*R*^*2*^ = .163, *R*_*adj*_^*2*^ = .162) of the variance in functional connectivity of novel validation samples, while the model accounted for 16.5% (*R*^*2*^ = .165, *R*_*adj*_^*2*^ = .164) of the variance in functional connectivity when predicting the data for all connections (null model permutation *p* = .012). These cross-validation results indicate that over-fitting is minimal. Finally, by investigating the coefficients for each of the independent variables, the pattern of results seen in the multiple linear regression model can be confirmed. For diffusion measures, mean first passage time had a coefficient of -.037, communicability had a coefficient of .018, and shortest path length had a coefficient of -.025. Note that the negative relationship between structure and function for mean first passage time and shortest path length was expected, as a higher value for these structural measures indicates weaker connectivity, while the positive relationship was expected for communicability as higher values indicate stronger connectivity. These coefficients support what was observed for the multiple linear regression, indicating that the effects of the diffusion model-based measures were greater in combined magnitude than that of shortest path length.

#### Principal components analysis

The principal components analysis for functional connectivity, mean first passage time, communicability, and shortest path length shown in Figure 3 demonstrates the unique component space occupied by each measure. This analysis was conducted using the *prcomp* function of the core *stats* library in *R*. To aid interpretation of the principal component loadings of each variable, mean first passage time and shortest path length were multiplied by -1 so that larger values indicate better connectivity for all measures. In particular, Principal Component 1 seems to be sensitive to the variance in common between functional connectivity and the graph theory measures, as these all have loadings in the same direction (Figure 3A and Table 4). Conversely, functional connectivity loads strongly onto Principal Component 2, while the graph theory measures load weakly and in the opposite direction, suggesting that this component identifies variance in functional connectivity that is not well accounted for by the graph theory measures (Figure 3A and Table 4). Finally, functional connectivity and shortest path length load very weakly onto Principal Component 3, while the loadings for mean first passage time and communicability are strong and in opposite directions, suggesting that this component speaks to the unique position in the component space of these diffusion model measures (Figure 3B and Table 4).

**Table 4.** AAL principal components analysis loadings for all 3 principal components. Variables considered were functional connectivity (FC), mean first passage time (MFPT; log transformed and multiplied by -1 so that more positive values indicate better connectivity), communicability (COM; log transformed), and shortest path length (SPL; log transformed and multiplied by -1 as with MFPT).

| Variable | PC1 | PC2 | PC3 |
| --- | --- | --- | --- |
| FC | -.359 | -.926 | .112 |
| -1*log(MFPT) | -.529 | .113 | -.671 |
| log(COM) | -.501 | .274 | .728 |
| -1*log(SPL) | -.583 | .232 | -.086 |

**Figure 2.**
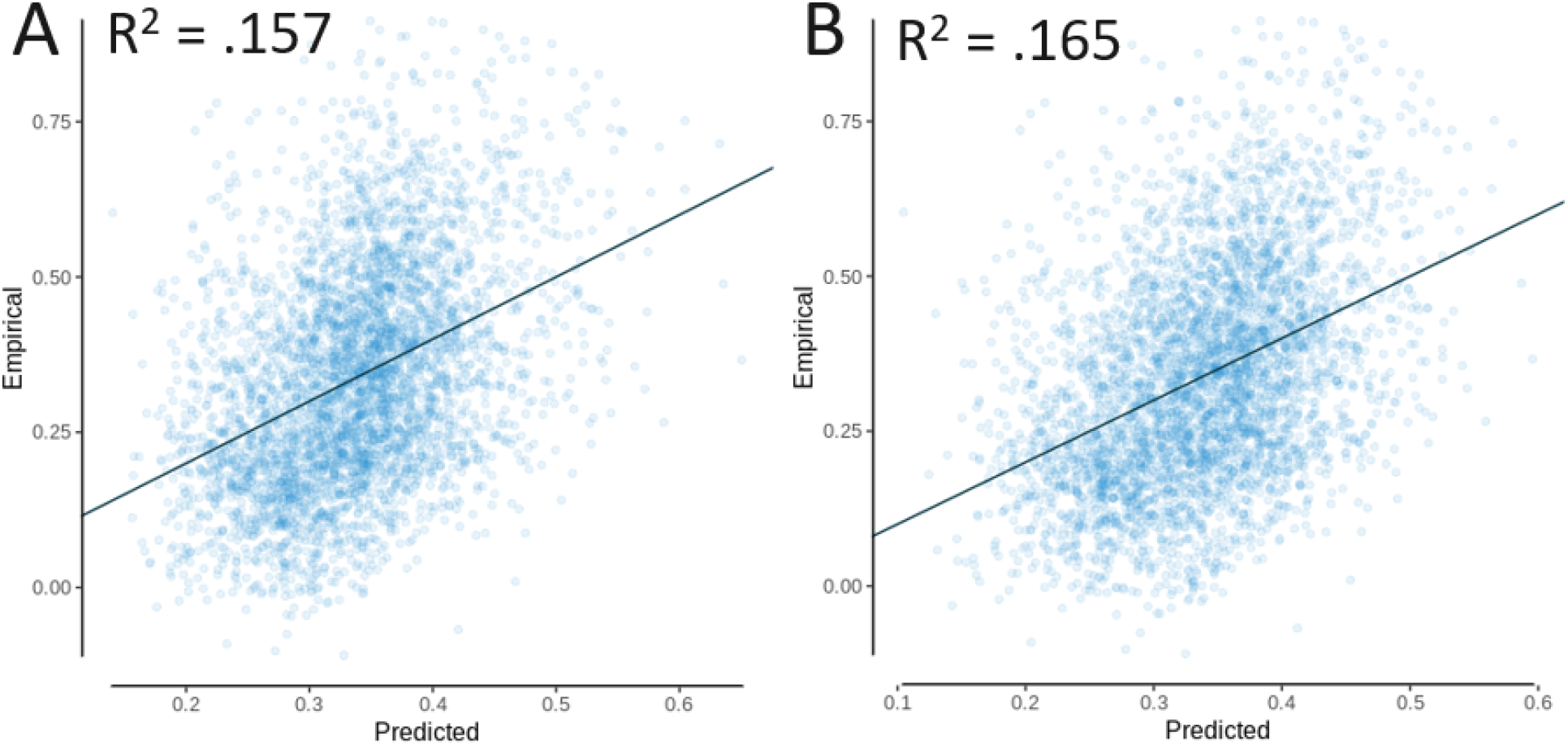
AAL multiple linear regression models with empirical functional connectivity as the as a function of the predicted functional connectivity, for **(A)** model 1 and **(B)** model 2.

**Figure 3.**
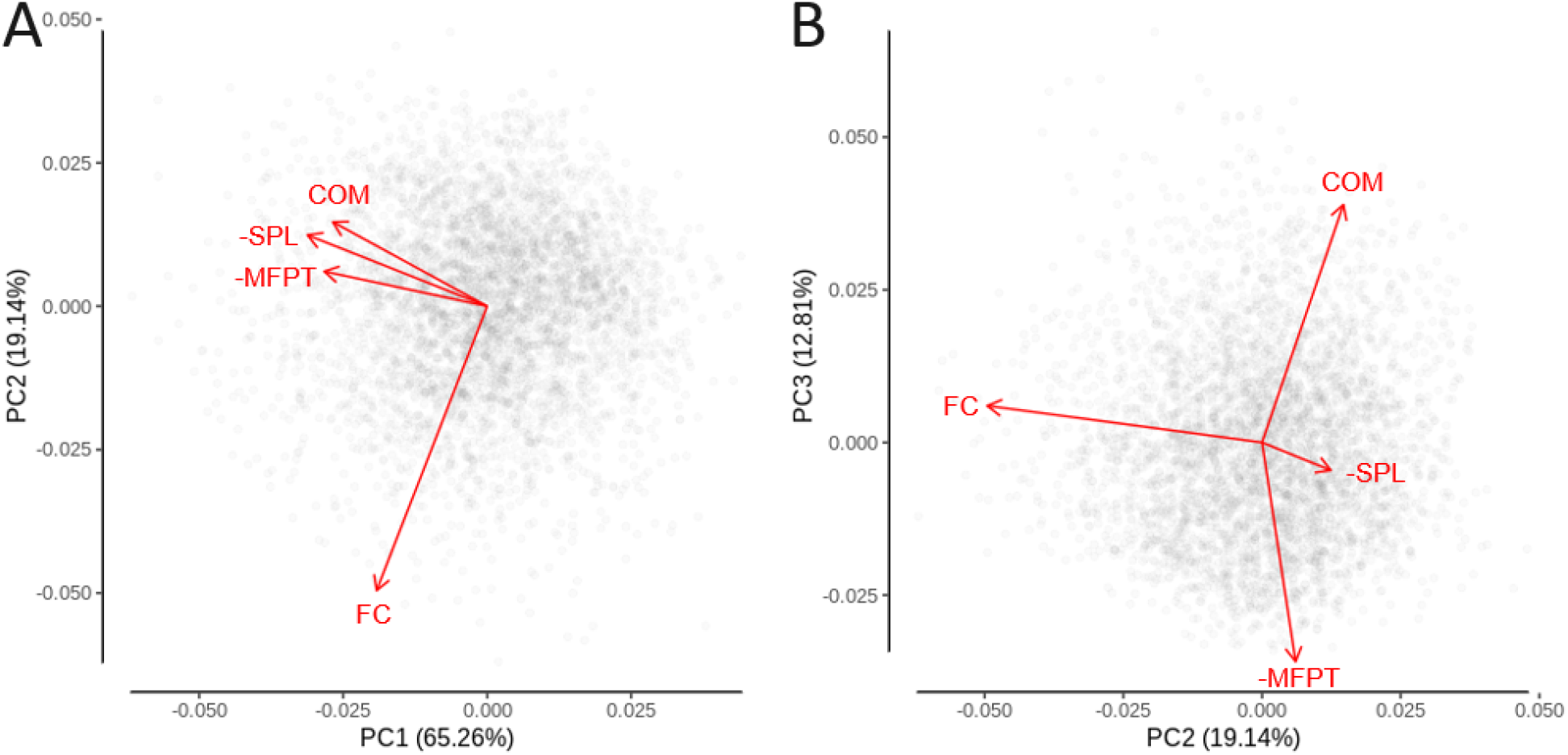
AAL principal components analysis with data points and variable loadings as vectors. Variables considered were functional connectivity (FC), mean first passage time (MFPT; log transformed and multiplied by -1 so that more positive values indicate better connectivity), communicability (COM; log transformed), and shortest path length (SPL; log transformed and multiplied by -1 as with MFPT).

### Brainnetome

#### Linear regression models

Using the Brainnetome atlas, there was again an inverse relationship between mean first passage time and functional connectivity, a positive relationship between communicability and functional connectivity, and an inverse relationship between shortest path length and functional connectivity. Again, the magnitude of the structure-function relationship for each of these measures was relatively comparable (see Table 5 and Figure 4).

**Table 5.**
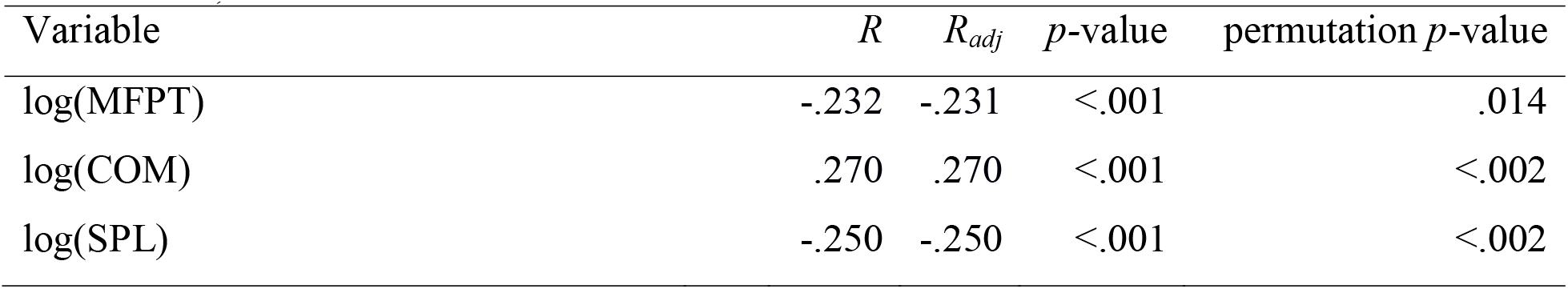
Brainnetome linear regression analyses. Dependent variable was functional connectivity, and independent variables considered were mean first passage time (MFPT; log transformed), communicability (COM; log transformed), and shortest path length (SPL; log transformed).

**Figure 4.**
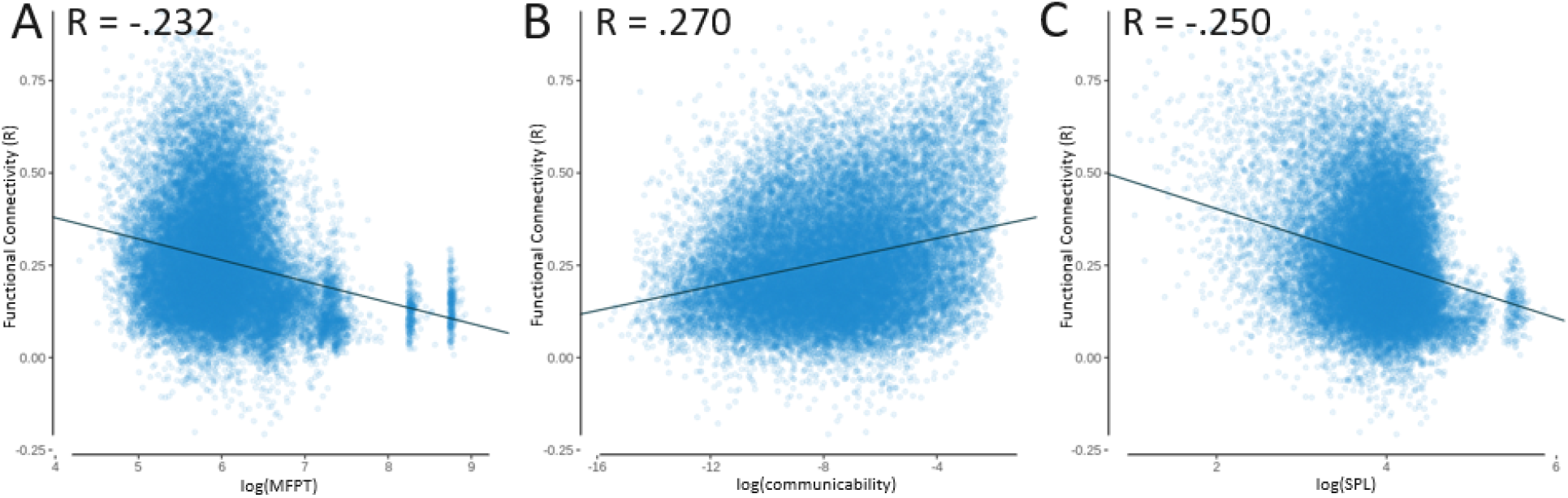
Brainnetome linear regression models with functional connectivity as the independent variable and dependent variables of: **(A)** mean first passage time (MFPT; log transformed), *R*_*adj*_ = -.231; **(B)** communicability (COM; log transformed), *R*_*adj*_ = .270; **(C)** and shortest path length (SPL; log transformed), *R*_*adj*_ = -.250. With outlier clusters removed from **(A)** by excluding cases with log(MFPT) > 8 and **(C)** by excluding cases with log(SPL) > 5.33 the correlation remains significant, with *R =* -.207 for log(MFPT) and *R* = -.224 for log(SPL). These outlier clusters are due to more isolated regions of the atlas that take more steps to reach than most other regions.

#### Multiple linear regression models

Multiple linear regression models were also investigated for the Brainnetome atlas, demonstrating that the semi-partial correlation for shortest path length was much less than the combined magnitude for the diffusion-based measures of mean first passage time and communicability (see Tables 6 through 8 and Figure 5).

**Table 6.** Brainnetome independent variable correlation matrix. Variables include mean first passage time (MFPT), communicability (COM), and shortest path length (SPL).

| Variable | MFPT | COM | SPL |
| --- | --- | --- | --- |
| MFPT | 1 | -.087 | .824 |
| COM | -.087 | 1 | -.342 |
| SPL | .824 | -.342 | 1 |

**Figure 5.**
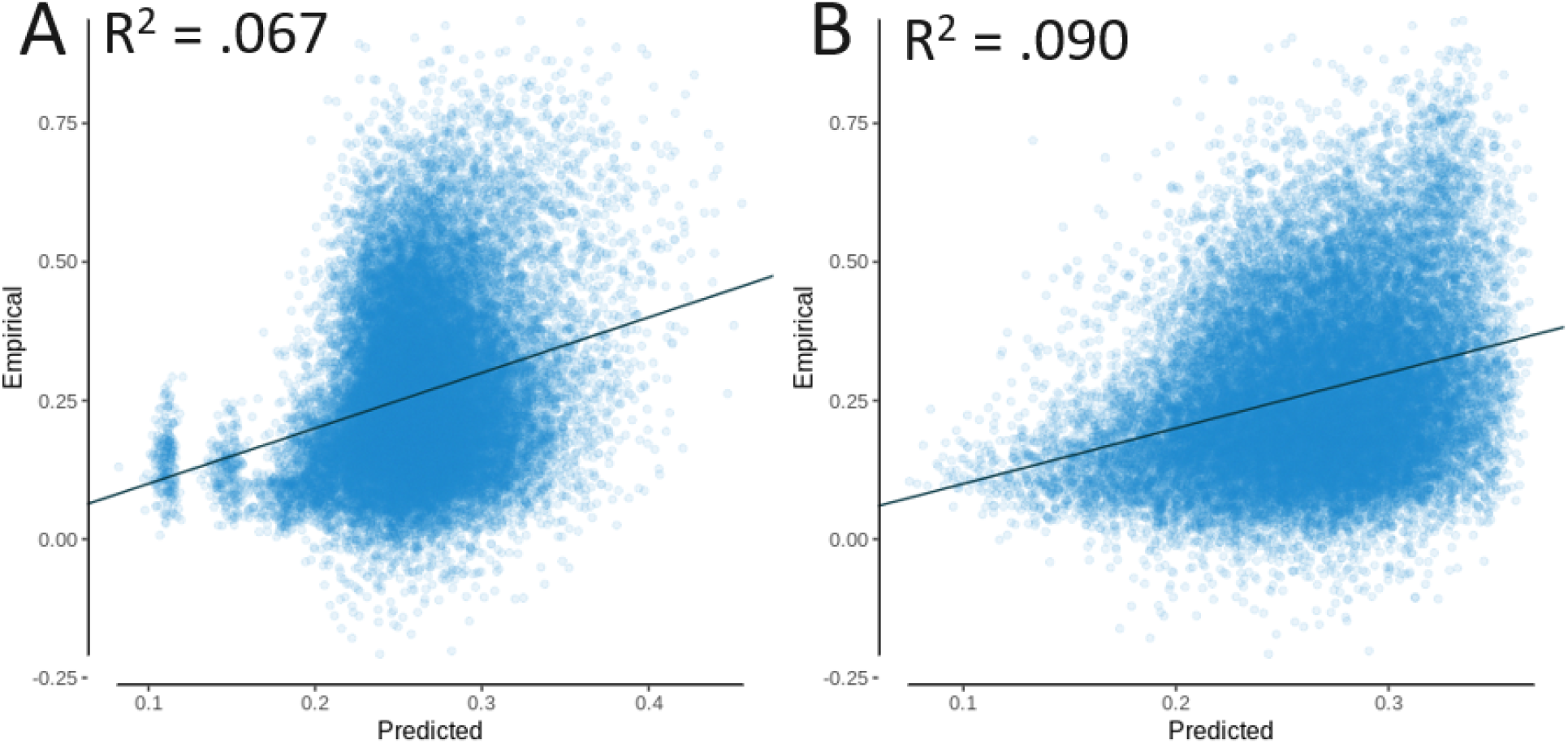
Brainnetome multiple linear regression models with empirical functional connectivity as a function of the predicted functional connectivity, for **(A)** model 1 and **(B)** model 2.

#### Partial least squares regression

The partial least squares regression again demonstrated that the combined magnitude of the coefficients for the diffusion model measures of mean first passage time and communicability were much greater than the shortest path routing model measure of shortest path length (see Table 9).

**Table 7.** Brainnetome multiple linear model 1, with dependent variable functional connectivity. *R*^*2*^ = .067, *R*_*adj*_^*2*^ = .067 (null model permutation *p* < .002). Abbreviations include semi-partial correlation (SPC) and variance inflation factor (VIF).

| Effects | SPC | Estimate | Std. Error | *t*-value | *p*-value | VIF |
| --- | --- | --- | --- | --- | --- | --- |
| Intercept |  | .614 | .008 | 73.783 | <.001 |  |
| log(MFPT) | -.067 | -.025 | .002 | -12.024 | <.001 | 2.275 |
| log(SPL) | -.116 | -.052 | .002 | -20.796 | <.001 | 2.275 |

**Table 8.** Brainnetome multiple linear model 2, with dependent variable functional connectivity. *R*^*2*^ = .090, *R*_*adj*_^*2*^ = .090 (null model permutation *p* < .002). Abbreviations include semi-partial correlation (SPC) and variance inflation factor (VIF).

| Effects | SPC | Estimate | Std. Error | *t*-value | *p*-value | VIF |
| --- | --- | --- | --- | --- | --- | --- |
| Intercept |  | .558 | .008 | 65.877 | <.001 |  |
| log(MFPT) | -.112 | -.044 | .002 | -20.383 | <.001 | 2.534 |
| log(COM) | .152 | .015 | $5.442 \times 10^{-4}$ | 27.643 | <.001 | 2.702 |
| log(SPL) | .034 | .022 | .004 | 6.107 | <.001 | 4.969 |

**Table 9.** Brainnetome partial least squares (PLS) regression. Dependent variable was functional connectivity, and independent variables considered were mean first passage time (MFPT; log transformed), communicability (COM; log transformed), and shortest path length (SPL; log transformed). Variance accounted for was *R*^*2*^ = .087, *R*_*adj*_^*2*^ = .087, null model permutation *p* < .002.

| Variable | Coefficient |
| --- | --- |
| log(MFPT) | -.015 |
| log(COM) | .035 |
| log(SPL) | -.003 |

#### Principal components analysis

The principal components analysis was again conducted, but this time for the Brainnetome atlas. This analysis replicated the pattern of results seen for the AAL atlas (see Figure 6 and Table 10).

**Figure 6.**
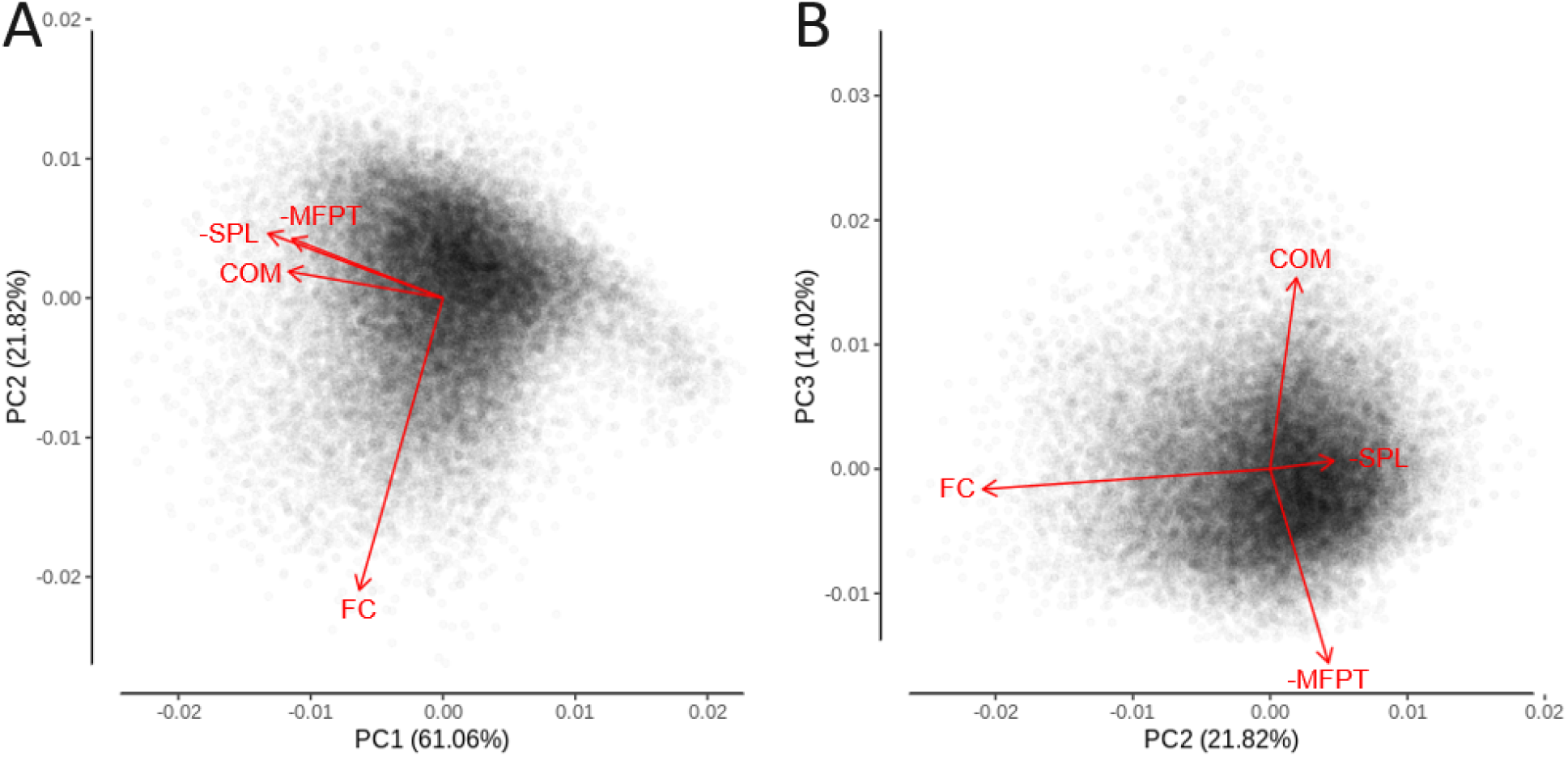
Brainnetome principal components analysis with data points and variable loadings as vectors. Variables considered were functional connectivity (FC), mean first passage time (MFPT; log transformed and multiplied by -1 so that more positive values indicate better connectivity), communicability (COM; log transformed), and shortest path length (SPL; log transformed and multiplied by -1 as with MFPT).

**Table 10.** Brainnetome principal components analysis loadings for all 3 principal components. Variables considered were functional connectivity (FC), mean first passage time (MFPT; log transformed and multiplied by -1 so that more positive values indicate better connectivity), communicability (COM; log transformed), and shortest path length (SPL; log transformed and multiplied by -1 as with MFPT).

| Variable | PC1 | PC2 | PC3 |
| --- | --- | --- | --- |
| FC | -.288 | -.954 | -.074 |
| -1*log(MFPT) | -.520 | .194 | -.710 |
| log(COM) | -.532 | .087 | .700 |
| -1*log(SPL) | -.603 | .211 | .030 |

### Supplementary Analyses

#### Individual-level analyses

Individual-level analyses were performed that mirrored the mean-level analyses, demonstrating that the variance accounted for was reduced but significant in all cases, with the same pattern of results as in the mean-level analyses (see Supplementary Tables 1 through 10). These analyses utilized the upper triangle of each connectivity matrix, reshaped to a single dimensional array for each individual and then concatenated across all individuals into a single large array for each of the mean first passage time, communicability, shortest path length, and functional connectivity measures. These arrays were used as variables in the statistical analyses in the same way as for the mean-level data.

#### Split-half analyses

Half of the data was used as a training set to train the models and the other half was used as a test set to test the models on novel data in order to demonstrate the predictability on out-of-sample data. In all cases the test sets were predicted with comparable accuracy (see Supplementary Tables 11 through 18).

#### PCA null models

Null models were used to produce a null distribution for loadings of each of the 4 variables on the 3 Principal Components, and demonstrated that there was a significant distance between the null and empirical PC loadings (see Supplementary Tables 19 and 20).

## Discussion

This investigation and comparison of graph theory structural connectivity measures based on two different theories of how information passes from one region to another in the brain has highlighted the importance of the diffusion model relative to the more straightforward but less biologically plausible shortest path routing model. Diffusion measures of mean first passage time and communicability as well as the shortest path routing measure of shortest path length were calculated from the brain structural connectivity, and these measures were related to functional connectivity. In isolation, each of these measures were comparable in the level to which they were related to functional connectivity. When analysed as hybrid models with each of these measures included, more variance was accounted for than any measure on its own, supporting past research suggesting that a hybrid/balance of both the diffusion and shortest path routing models is appropriate for describing the structure-function relationship (Goñi et al., 2013, 2014). However, when analysed together in multiple linear regression, partial least squares regression, and principal components analysis, it was clear that the diffusion model and the ideas it can express capture more of the variance in functional connectivity than shortest path routing, suggesting that the diffusion model may be closer to describing how information travels from one region to another in the structural connectivity network to produce the observed patterns of functional connectivity. These findings suggest that shortest path routing may be somewhat redundant when diffusion models are able to travel along similarly efficient routes in the brain, and that diffusion models are able to additionally tap into aspects of functional connectivity that are not considered by shortest path routing. These results are not surprising, given that diffusion models are more biologically plausible than shortest path routing as there is no evidence to suggest that regions have global network knowledge about what path would be the shortest (Avena-Koenigsberger et al., 2019; Seguin et al., 2018, 2022; Zamani Esfahlani et al., 2022). Past research has demonstrated that both diffusion models and shortest path routing models are important frameworks for understanding the architecture of brain networks (Goñi et al., 2013, 2014), but this work has taken an important next step in distinguishing the greater relative ability for the diffusion measures to accurately predict function from the underlying structural connectivity.

### Limitations and Future Directions

While diffusion and shortest path routing models are the most commonly discussed in network neuroscience, other models have been suggested that may add a unique perspective on how information travels in the brain. One such proposed theory for how information may transfer is *greedy navigation*, in which the Euclidean (three dimensional) or geodesic (two-dimensional flattened surface of cortex) distance between regions is used to travel through the network to whichever region is closest to the target region. This model has been investigated in simulated networks, which have shown that greedy navigation is able to successfully send information between regions without getting stuck (i.e., ending up at a node that is closer than all neighbours to the destination but unconnected to the destination) when there is a balance between clustered connections of proximal nodes and a power-law like distribution whereby clusters are connected by a small number of highly connected hub nodes (Boguñá et al., 2009). The pattern of navigation under these conditions is such that the path tends to travel to a nearby hub node, travel a long distance to another hub node, then travel to a low degree node in a cluster close to the destination. This pattern has also been demonstrated in the macaque brain network (Harriger et al., 2012) and in *C. elegans* (Towlson et al., 2013). Research using this theory in the human brain has demonstrated prediction of functional connectivity from structural connectivity (Seguin et al., 2018, 2020). Future research should implement a model of *greedy navigation* to replicate that the path length of a greedy navigator predicts functional connectivity, and additionally investigate whether the pattern of paths resembles that seen in simulated models and animal models.

Internet and computer analogies have also been proposed for application to brain networks, with encouraging results (Graham & Rockmore, 2011; Mišić et al., 2014). Information flow in computer networks (such as the internet) is limited by bandwidth, which represents the amount of information that can travel in a certain amount of time (e.g., bits/s). Information can be transmitted via packet switching, which breaks messages into packets that are labeled with the intended destination. These packets traverse the network efficiently by utilizing connections at separate times and buffering in a node if a connection is full, with the downside that if the node buffer is full then the information is lost. The structural connectivity of the macaque has been used to simulate a message-switched variant of this model of information flow, showing that compared to other networks there was more message loss, lower throughput, but faster transit times (Mišić et al., 2014). This suggests that under the model assumptions speed would be optimized in the macaque brain over maintaining the integrity of each individual signal. Future research should work on also applying a packet switching simulation model using the human brain to uncover whether similar signatures of information flow are seen between human and macaque networks, and whether the time taken for information to travel between regions is predictive of functional connectivity measures.

As another example of further work to be done in the field, brain network analysis of the cat brain has found that regions with similar connectivity profiles (connect to the same or similar regions) tended to correspond with groups of regions performing similar tasks (Zamora-López et al., 2010). One measure of the similarity between connectivity profiles of regions is *cosine similarity*, which has been used to demonstrate that functionally similar regions in the macaque brain also had high cosine similarity, and that regions with high cosine similarity were also likely to be connected and located close together (Song et al., 2014). This measure has recently been investigated along with many other measures in the context of the human brain (Zamani Esfahlani et al., 2022), and more research should be done to investigate whether human brain regions with similar connectivity profiles are more likely to be connected, to what extent this pattern differs between functionally similar clusters and core hub regions, and to what extent the structural cosine similarity is able to predict the functional connectivity and functional cosine similarity.

Finally, a recent line of research has investigated the potential of hybrid models of information transfer that allow for regional heterogeneity in the navigation strategy applied. This is an extension of research that has shown differences in structure-function coupling when looking at unimodal vs. transmodal regions, and development of these patterns with age (Baum et al., 2020). These models allow for different regions to apply unique strategies of information transfer, and have shown promising findings (Avena-Koenigsberger et al., 2019; Vázquez-Rodríguez et al., 2019; Zamani Esfahlani et al., 2022). This is a promising line of research that we expect will continue improving our understanding of how information is propagated throughout the structural connectivity network of the brain.

## Conclusion

Although the models described here do not claim to be a fully accurate and comprehensive description of how information is transferred in the brain, especially considering that the nature of the structure-function relationship will inevitably vary as the scale of interest goes from the macroscale to the microscale, by examining the diffusion and shortest path routing models together, this research has demonstrated that diffusion models are better suited to describing the relationship between structural and functional connectivity at the macroscale. In the future, alternative models discussed here could be examined together, in order to contribute to a fuller picture of which aspects of these theoretical models are able to best approximate the ground truth of information transfer in the human brain network.

## Acknowledgements

This research was supported by the Natural Sciences and Engineering Research Council of Canada through Alexander Graham Bell Canada Graduate Scholarships to Josh Neudorf and Shaylyn Kress, and Discovery Grant 18968-2013-25 to the senior author Ron Borowsky. The authors affirm that there are no conflicts of interest to disclose.

Data were provided [in part] by the Human Connectome Project, WU-Minn Consortium (Principal Investigators: David Van Essen and Kamil Ugurbil; 1U54MH091657) funded by the 16 NIH Institutes and Centers that support the NIH Blueprint for Neuroscience Research; and by the McDonnell Center for Systems Neuroscience at Washington University.

## Declarations

### Availability of data and material

All data is made publicly available by the Human Connectome Project (Van Essen et al., 2013).

### Code availability

The code used to apply the equations is available upon reasonable request.

### Ethics approval, consent to participate, and consent for publication

Please see Van Essen et al. (2013) for a description of ethics and consent.

## Supplementary Materials

**Supplementary Table 1.**
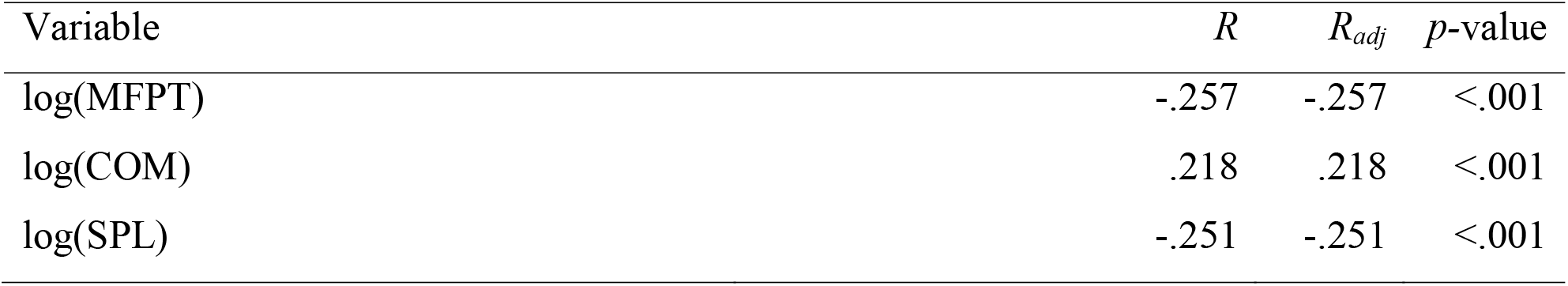
AAL individual-level linear regression analyses. Dependent variable was functional connectivity, and independent variables considered were mean first passage time (MFPT; log transformed), communicability (COM; log transformed), and shortest path length (SPL; log transformed).

**Supplementary Table 2.** AAL Individual-level multiple linear model 1, with dependent variable functional connectivity. *R*^*2*^ = .072, *R*_*adj*_^*2*^ = .072. Abbreviations include semi-partial correlation (SPC) and variance inflation factor (VIF).

| Effects | SPC | Estimate | Std. Error | $t$-value | $p$-value | VIF |
| --- | --- | --- | --- | --- | --- | --- |
| Intercept | | .772 | $9.377 \times 10^{-4}$ | 822.769 | <.001 | |
| log(MFPT) | -.092 | -.056 | $2.931 \times 10^{-4}$ | -191.660 | <.001 | 2.844 |
| log(SPL) | -.075 | -.047 | $3.059 \times 10^{-4}$ | -155.079 | <.001 | 2.844 |

**Supplementary Table 3.** AAL Individual-level multiple linear model 2, with dependent variable functional connectivity. *R*^*2*^ = .078, *R*_*adj*_^*2*^ = .078. Abbreviations include semi-partial correlation (SPC) and variance inflation factor (VIF).

| Effects | SPC | Estimate | Std. Error | $t$-value | $p$-value | VIF |
| --- | --- | --- | --- | --- | --- | --- |
| Intercept | | .771 | $9.347 \times 10^{-4}$ | 825.186 | <.001 | |
| log(MFPT) | -.103 | -.064 | $2.958 \times 10^{-4}$ | -215.179 | <.001 | 2.915 |
| log(COM) | .077 | .011 | $6.906 \times 10^{-5}$ | 161.222 | <.001 | 1.960 |
| log(SPL) | -.017 | -.013 | $3.724 \times 10^{-4}$ | -34.832 | <.001 | 4.242 |

**Supplementary Table 4.** AAL individual-level partial least squares (PLS) regression. Dependent variable was functional connectivity, and independent variables considered were mean first passage time (MFPT; log transformed), communicability (COM; log transformed), and shortest path length (SPL; log transformed). Variance accounted for was *R*^*2*^ = .077, *R*_*adj*_^*2*^ = .077.

| Variable | Coefficient |
| --- | --- |
| log(MFPT) | -.039 |
| log(COM) | .019 |
| log(SPL) | -.017 |

**Supplementary Table 5.** AAL individual-level principal components analysis loadings for all 3 principal components. Variables considered were functional connectivity (FC), mean first passage time (MFPT; log transformed and multiplied by -1 so that more positive values indicate better connectivity), communicability (COM; log transformed), and shortest path length (SPL; log transformed and multiplied by -1 as with MFPT).

| Variable | PC1 | PC2 | PC3 |
| --- | --- | --- | --- |
| FC | -.278 | -.960 | .035 |
| -1*log(MFPT) | -.552 | .126 | -.609 |
| log(COM) | -.511 | .170 | .783 |
| -1*log(SPL) | -.597 | .185 | -.124 |

**Supplementary Table 6.**
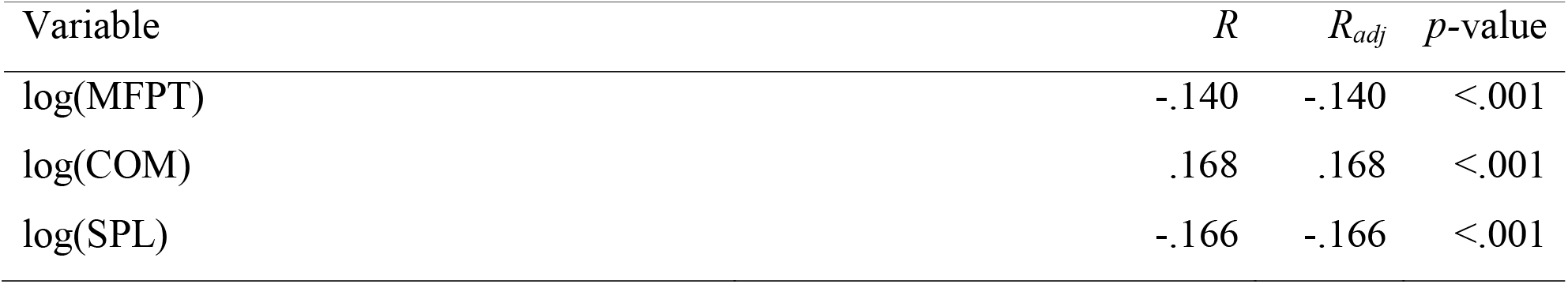
Brainnetome individual-level linear regression analyses. Dependent variable was functional connectivity, and independent variables considered were mean first passage time (MFPT; log transformed), communicability (COM; log transformed), and shortest path length (SPL; log transformed).

**Supplementary Table 7.**
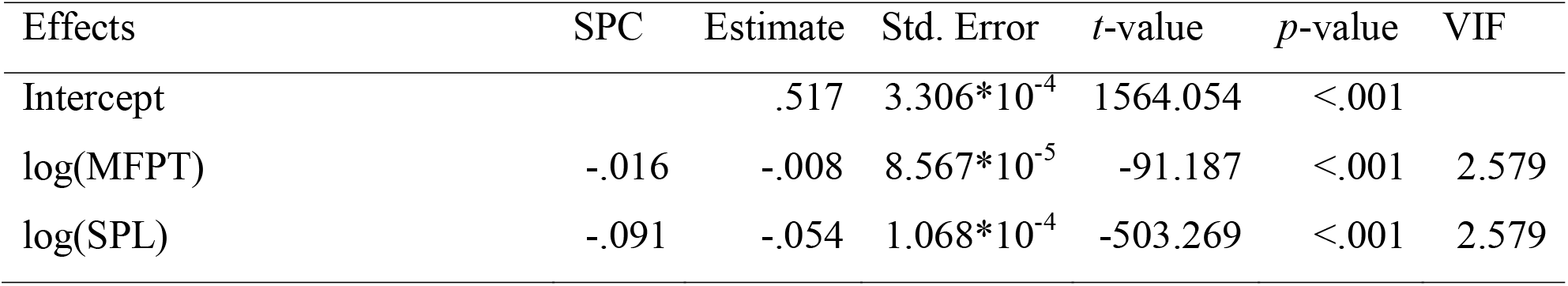
Brainnetome individual-level multiple linear model 1, with dependent variable functional connectivity. *R*^*2*^ = .028, *R*_*adj*_^*2*^ = .028. Abbreviations include semi-partial correlation (SPC) and variance inflation factor (VIF).

| Effects | SPC | Estimate | Std. Error | *t*-value | *p*-value | VIF |
| --- | --- | --- | --- | --- | --- | --- |
| Intercept | | .517 | $3.306 \times 10^{-4}$ | 1564.054 | <.001 | |
| log(MFPT) | -.016 | -.008 | $8.567 \times 10^{-5}$ | -91.187 | <.001 | 2.579 |
| log(SPL) | -.091 | -.054 | $1.068 \times 10^{-4}$ | -503.269 | <.001 | 2.579 |

**Supplementary Table 8.** Brainnetome individual-level multiple linear model 2, with dependent variable functional connectivity. *R*^*2*^ = .034, *R*_*adj*_^*2*^ = .034. Abbreviations include semi-partial correlation (SPC) and variance inflation factor (VIF).

| Effects | SPC | Estimate | Std. Error | *t*-value | *p*-value | VIF |
| --- | --- | --- | --- | --- | --- | --- |
| Intercept | | .500 | $3.317 \times 10^{-4}$ | 1505.702 | <.001 | |
| log(MFPT) | -.032 | -.016 | $8.717 \times 10^{-5}$ | -180.304 | <.001 | 2.689 |
| log(COM) | .081 | .008 | $1.814 \times 10^{-5}$ | 449.561 | <.001 | 1.896 |
| log(SPL) | -.025 | -.018 | $1.322 \times 10^{-4}$ | -139.992 | <.001 | 3.978 |

**Supplementary Table 9.**
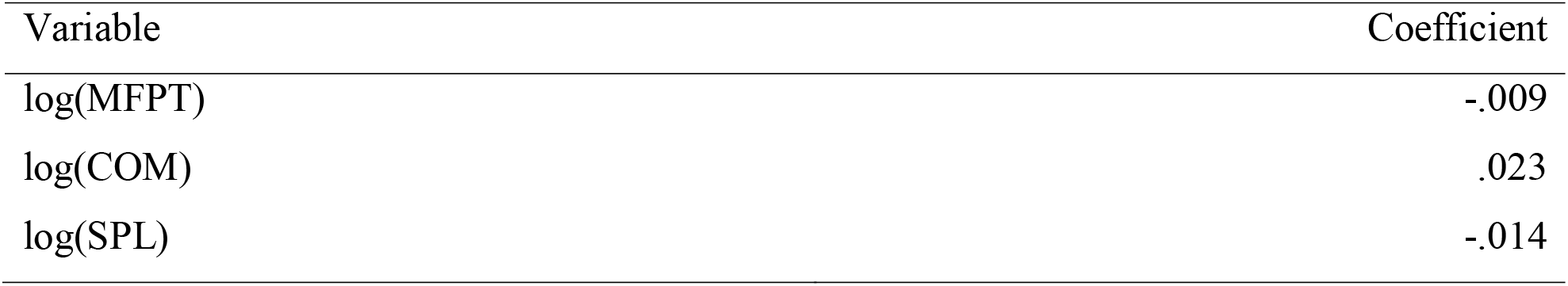
Brainnetome individual-level partial least squares (PLS) regression. Dependent variable was functional connectivity, and independent variables considered were mean first passage time (MFPT; log transformed), communicability (COM; log transformed), and shortest path length (SPL; log transformed). Variance accounted for was *R*^*2*^ = .034, *R*_*adj*_^*2*^ = .034.

**Supplementary Table 10.** Brainnetome individual-level principal components analysis loadings for all 3 principal components. Variables considered were functional connectivity (FC), mean first passage time (MFPT; log transformed and multiplied by -1 so that more positive values indicate better connectivity), communicability (COM; log transformed), and shortest path length (SPL; log transformed and multiplied by -1 as with MFPT).

| Variable | PC1 | PC2 | PC3 |
| --- | --- | --- | --- |
| FC | -.202 | -.977 | -.074 |
| -1*log(MFPT) | -.556 | .158 | -.609 |
| log(COM) | -.520 | .046 | .785 |
| -1*log(SPL) | -.617 | .139 | -.088 |

**Supplementary Table 11.**
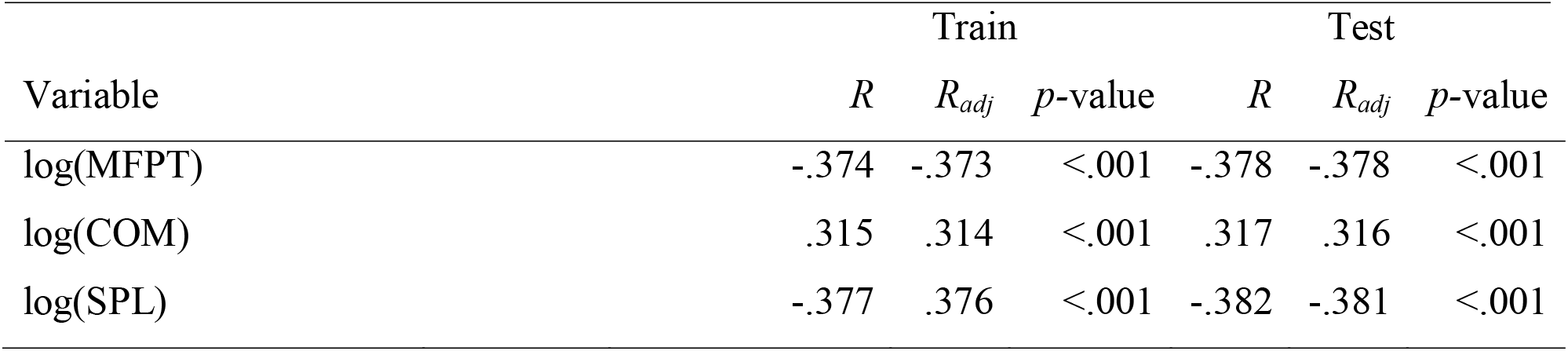
AAL split-half linear regression analyses. Model was trained on a training set and then tested on a testing set. Dependent variable was functional connectivity, and independent variables considered were mean first passage time (MFPT; log transformed), communicability (COM; log transformed), and shortest path length (SPL; log transformed).

**Supplementary Table 12.** AAL split-half multiple linear model 1. Model was trained on a training set and then tested on a testing set, with dependent variable functional connectivity. Variance accounted for in training set was *R*^*2*^ = .155, *R*_*adj*_^*2*^ = .155, and in testing set was *R*^*2*^ = .159, *R*_*adj*_^*2*^ = .159. Abbreviations include semi-partial correlation (SPC) and variance inflation factor (VIF).

| Effects | SPC | Estimate | Std. Error | t-value | p-value | VIF |
| --- | --- | --- | --- | --- | --- | --- |
| Intercept |  | .838 | .023 | 36.235 | <.001 |  |
| log(MFPT) | -.115 | -.059 | .007 | -7.947 | <.001 | 2.946 |
| log(SPL) | -.126 | -.066 | .008 | -8.657 | <.001 | 2.946 |

**Supplementary Table 13.**
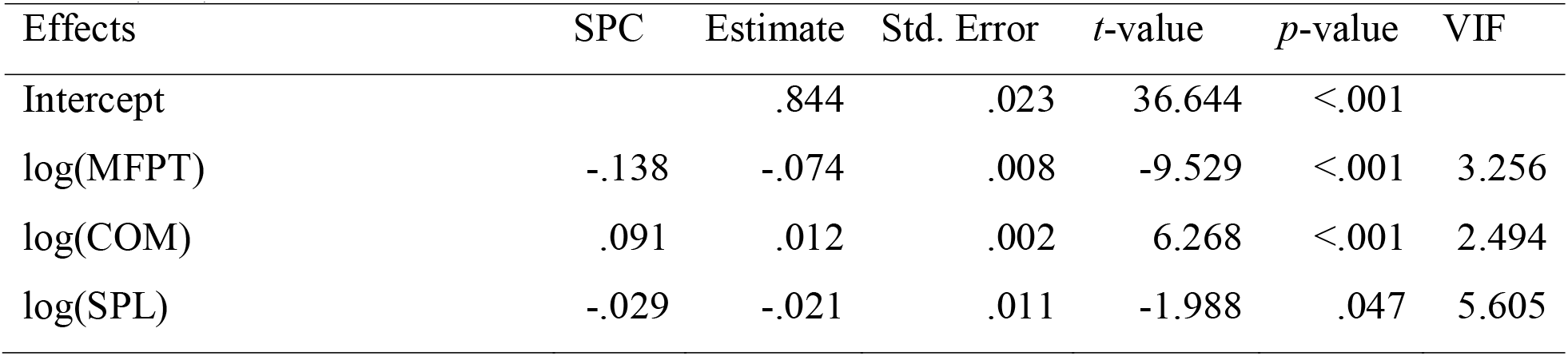
AAL split-half multiple linear model 2. Model was trained on a training set and then tested on a testing set, with dependent variable functional connectivity. Variance accounted for in training set was *R*^*2*^ = .164, *R*_*adj*_^*2*^ = .163, and in testing set was *R*^*2*^ = .167, *R*_*adj*_^*2*^ = .166. Abbreviations include semi-partial correlation (SPC) and variance inflation factor (VIF).

**Supplementary Table 14.** AAL split-half partial least squares (PLS) regression. Model was trained on a training set and then tested on a testing set. Dependent variable was functional connectivity, and independent variables considered were mean first passage time (MFPT; log transformed), communicability (COM; log transformed), and shortest path length (SPL; log transformed). Variance accounted for in training set was *R*^*2*^ = .163, *R*_*adj*_^*2*^ = .162, and in testing set was *R*^*2*^ = .166, *R*_*adj*_^*2*^ = .166.

| Variable | Coefficient |
| --- | --- |
| log(MFPT) | -.037 |
| log(COM) | .018 |
| log(SPL) | -.024 |

**Supplementary Table 15.** Brainnetome split-half linear regression analyses. Model was trained on a training set and then tested on a testing set. Dependent variable was functional connectivity, and independent variables considered were mean first passage time (MFPT; log transformed), communicability (COM; log transformed), and shortest path length (SPL; log transformed).

| Variable | Train |  |  | Test |  |  |
| --- | --- | --- | --- | --- | --- | --- |
|  | *R* | *R<sub>adj</sub>* | *p*-value | *R* | *R<sub>adj</sub>* | *p*-value |
| log(MFPT) | -.230 | -.230 | <.001 | -.233 | -.233 | <.001 |
| log(COM) | .268 | .268 | <.001 | .272 | .272 | <.001 |
| log(SPL) | -.248 | -.248 | <.001 | -.252 | -.251 | <.001 |

**Supplementary Table 16.** Brainnetome split-half multiple linear model 1. Model was trained on a training set and then tested on a testing set, with dependent variable functional connectivity. Variance accounted for in training set was *R*^*2*^ = .066, *R*_*adj*_^*2*^ = .066, and in testing set was *R*^*2*^ = .068, *R*_*adj*_^*2*^ = .068. Abbreviations include semi-partial correlation (SPC) and variance inflation factor (VIF).

| Effects | SPC | Estimate | Std. Error | *t*-value | *p*-value | VIF |
| --- | --- | --- | --- | --- | --- | --- |
| Intercept |  | .606 | .008 | 73.398 | <.001 |  |
| log(MFPT) | -.066 | -.024 | .002 | -11.805 | <.001 | 2.295 |
| log(SPL) | -.115 | -.051 | .002 | -20.588 | <.001 | 2.295 |

**Supplementary Table 17.**
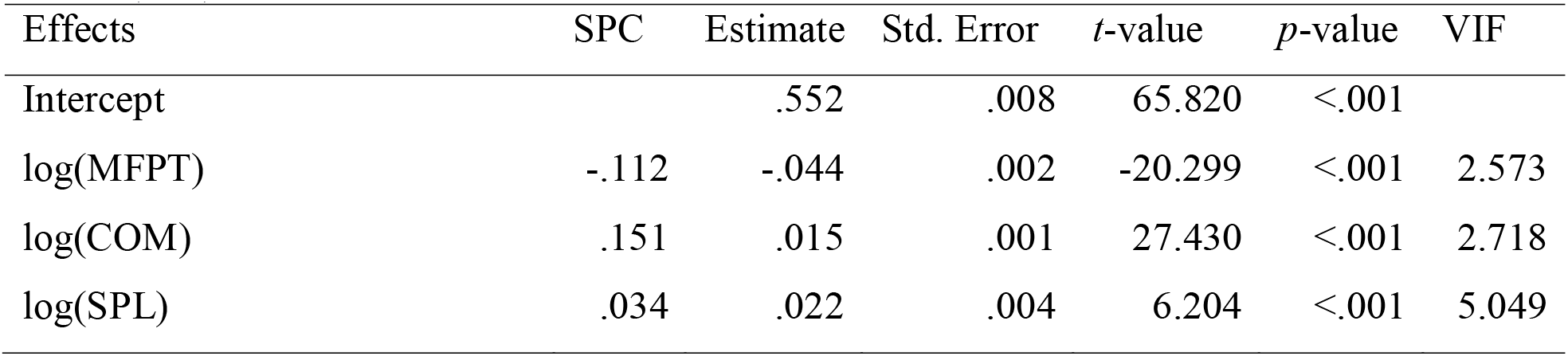
Brainnetome split-half multiple linear model 2. Model was trained on a training set and then tested on a testing set, with dependent variable functional connectivity. Variance accounted for in training set was *R*^*2*^ = .089, *R*_*adj*_^*2*^ = .089, and in testing set was *R*^*2*^ = .091, *R*_*adj*_^*2*^ = .091. Abbreviations include semi-partial correlation (SPC) and variance inflation factor (VIF).

| Effects | SPC | Estimate | Std. Error | *t*-value | *p*-value | VIF |
| --- | --- | --- | --- | --- | --- | --- |
| Intercept |  | .552 | .008 | 65.820 | <.001 |  |
| log(MFPT) | -.112 | -.044 | .002 | -20.299 | <.001 | 2.573 |
| log(COM) | .151 | .015 | .001 | 27.430 | <.001 | 2.718 |
| log(SPL) | .034 | .022 | .004 | 6.204 | <.001 | 5.049 |

**Supplementary Table 18.** Brainnetome split-half partial least squares (PLS) regression. Model was trained on a training set and then tested on a testing set. Dependent variable was functional connectivity, and independent variables considered were mean first passage time (MFPT; log transformed), communicability (COM; log transformed), and shortest path length (SPL; log transformed). Variance accounted for in training set was *R*^*2*^ = .086, *R*_*adj*_^*2*^ = .086 and in testing set was *R*^*2*^ = .088, *R*_*adj*_^*2*^ = .088.

| Variable | Coefficient |
| --- | --- |
| log(MFPT) | -.015 |
| log(COM) | .034 |
| log(SPL) | -.003 |

**Supplementary Table 19.** AAL null model mean distance from empirical PC vectors in PCA analysis, using a 3-dimensional distance calculation with the 3 PCs as dimensions. Bootstrapped 95% confidence intervals (95% CI) indicate that all measures were significantly different than the empirical PCs.

| Variable | Mean Distance | 95% CI |  |
| --- | --- | --- | --- |
| FC | .673 | .634 | .717 |
| -1*log(MFPT) | .915 | .881 | .946 |
| log(COM) | .838 | .804 | .870 |
| -1*log(SPL) | .980 | .948 | 1.010 |

**Supplementary Table 20.**
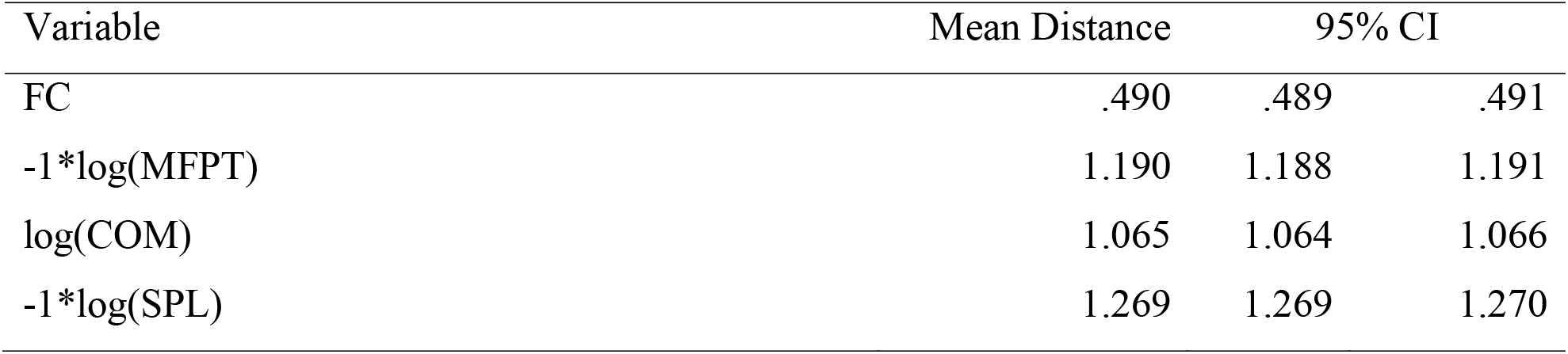
Brainnetome null model mean distance from empirical PC vectors in PCA analysis, using a 3-dimensional distance calculation with the 3 PCs as dimensions. Bootstrapped 95% confidence intervals (95% CI) indicate that all measures were significantly different than the empirical PCs.

